# The Symmetry and Asymmetry Behind Histone Folding Across Eukarya and Archaea

**DOI:** 10.1101/2022.10.15.512373

**Authors:** Haiqing Zhao, Hao Wu, Alex Guseman, Dulith Abeykoon, Christina M. Camara, Yamini Dalal, David Fushman, Garegin A. Papoian

## Abstract

Histones are the dominant proteins to compact and store DNA in both Eukarya and Archaea. For a long time, histones are observed to exist in the unit of dimers but diverge into different formats such as heterodimers in Eukarya or homodimers in Archaea. Here, by studying 11 types of histone proteins, both monomers and their dimeric complexes, using multiscale molecular dynamics (MD) simulations combined with NMR and circular dichroism experiments, we confirm the widely applied “folding upon binding” mechanism of histone structures. A histone dimer appears to form the longest *α*2 helices followed by other shorter helices and inter-molecular tertiary structures. We report an alternative conformation, namely, the inverted non-native dimer, which has a minimum free energy state. Protein sequence analysis indicates that the inverted conformation can be attributed to a hidden head-tail sequence symmetry underlying all histone proteins. This finding strongly support previously proposed histone evolution hypotheses. Finally, we separately used the MD-based AWSEM and AI-based AlphaFold-Multimer model to predict eukaryotic histone homodimer structures and performed extensive allatom MD simulations to examine their structural stabilities. Our results suggest that eukaryotic histones can also form stable homodimers, whereas their disordered tails— the structurally asymmetrical region—may tip the balance towards the formation of heterotypic dimers.

## INTRODUCTION

In Eukarya, histones are fundamental proteins for chromosome packaging. In nucleosome, the basic unit of chromosome, histones are assembled as four dimers, two H3/H4 heterodimers forming a tetramer and two H2A/H2B heterodimers separately^1^. Besides the canonical histones composing the majority of eukaryotic nucleosomes, histone variants and other histone-like proteins carry out specific functions in the nucleus^2,3^. In Archaea, histones are encoded to package and compact DNA into continuous hypernucleosomes^4,5^. Among many different histone oligomerization variations, dimer is the smallest unit reported in both Eukarya and Archaea, with regard to either structure or function. Nevertheless, in Eukarya histones exist as hetero-dimers with long terminal tails (Figure 1A), while in Archaea both homo- and hetero-dimer exist with no histone tails (Figure 1B).

**Figure 1:**
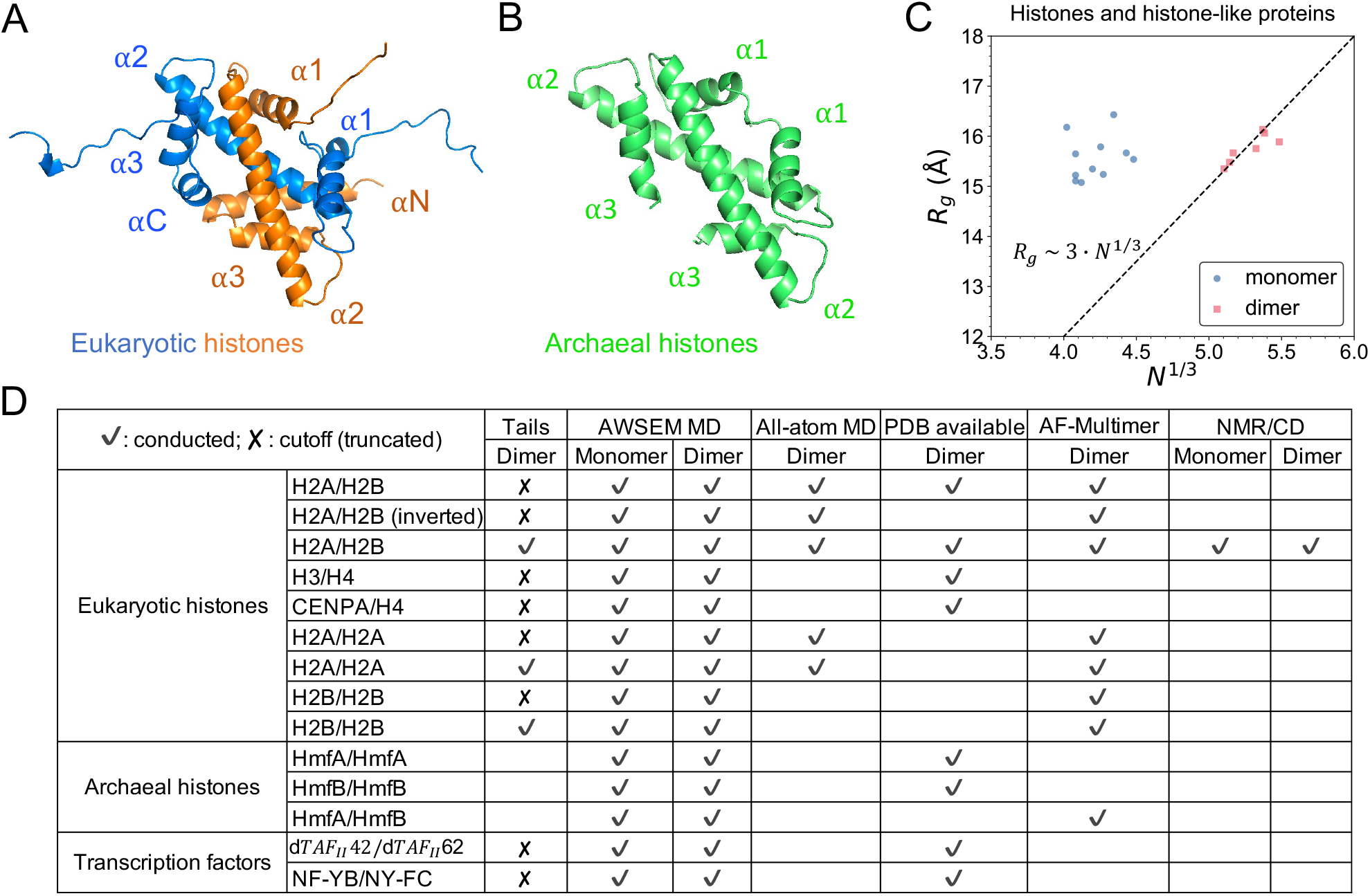
Structures of eukaryotic and archaeal histones and a summary table of studied systems. (A) The scaffold of eukaryotic histone dimer H2A/H2B (blue/orange, PDB: 1AOI) consists of three *α* helices in each protein, plus two terminal helices and long histone tails. (B) The structure of homotypic archaeal histone dimer (HmfB)_2_ (green, PDB: 1A7W) has three *α* helices each with no histone tails. (C) Polymer scaling fitting of the *R*_*g*_ and sequence length suggests that histone dimers act more as “monomeric” proteins. 7 histone(-like) dimer structures from PDB and their 11 monomers are included in this plot. The dashed line is the empirical relation of *R*_*g*_ and *N* from a survey study of 403 globular monomeric proteins^25^. (D) This table summarzies the 13 types of heterotypic and homotypic histone(-like) dimers that are studied in this work by different methodologies. A blank cell in Tails/PDB column means “non-existent” while in other columns it means “not conducted”.

Despite their function and sequence diversities across species, histones and histone-like proteins possess the same structural motif, known as histone-fold, which is comprised of three alpha helices connected by two loops^6^. Two histone-fold monomers assemble into a “handshake” pattern forming a dimer in an intertwined, head-to-tail manner^7^ (Figure 1A/B). A scaling analysis of the radius of gyration (*R*_*g*_) as a function of the protein size suggests that histone dimer acts as the minimum folded state, rather than its monomer (Figure 1C, supplemental session S1). Indeed, experimentally, it was known that eukaryotic histones fold and form complexes only in presence of their binding partner^8,9^. The stability of H2A/H2B dimers and (H3/H4)_2_ tetramers were studied by a series of denaturation experiments^10–13^. Karantza *et al*. reported that during the unfolding of either H2A/H2B or (H3/H4)_2_, individual folded monomers were not detectable, indicating a direct transition from one folded histone dimer to two unfolded monomers.

However, the molecular principles underlying this cooperative behavior have not been elucidated. In addition, whether or not these principles vary among eukaryotic histones, archaeal histones, and other histone-like structures remain unknown. Deeper insights into the folding dynamics of histone and histone-like proteins can not only shed light on their evolutionary basis and divergence, but also help understand their higher level structural organizations such as tetramer or octamer formation^14,15^, nucleosome interactome^16,17^, and chromosome organization^18–22^. In this work, we aim to build mechanistic understanding of histone folding at molecular detail and more broadly, to compare the folding mechanisms of histones across Eukarya and Archaea.

To address these questions, we first employed molecular dynamics (MD) simulations based on a coarse-grained protein force field, AWSEM^23^, which has been successfully applied to predict structures of proteins and their complexes^23,24^. Using AWSEM, we studied the protein foldings of seven different histones and four histone-like transcription factors (Figure 1D). We find that all studied histones or histone-like proteins are unstable as monomers. In agreement with the abovementioned prior experiments, our simulations show that two histone monomers cooperatively fold into a stable complex, revealing folding-upon-binding dynamics. We complemented our computational predictions by experimental investigations using Nuclear Magnetic Resonance (NMR) spectroscopy and circular dichroism (CD). The experimental data indicate that histones adopt an ordered structure only upon binding their partners at equimolar stoichiometry. All studied systems and used methodologies are summarized in Figure 1D.

Subsequent free energy (FE) calculations revealed an unexpected inverted non-native conformation of the histone dimer which occupies an FE minimum. The energy barrier between the native and non-native states was estimated around 8-9 kcal/mol. Further analysis of protein sequence alignment shows that our observation of non-native conformation is consistent with the well-known evolution hypotheses of histone-fold structural motif^26^, unveiling a hidden sequence symmetry in contemporary histones. Moreover, our simulations suggest that archaeal histones and other histone-like proteins including transcription factors also exhibit folding upon binding, with the abovementioned non-native dimer conformation potentially being a low-energy state.

Lastly, we applied the AWSEM model as well as the deep-learning structure prediction algorithm AlphaFold-Multimer (AF-Multimer)^27,28^ to predict the structure of hypothetical eukaryotic histone homodimer and the potential effects of histone tails. Structures obtained from these two independent methodologies were examined through extensive all-atom MD simulations. Together, these simulations suggest the important role of histone tails in determining the folding and formations of various eukaryotic histone dimers, which distinguishes eukaryotic histones from archaeal ones.

## METHODS

### MD Simulations and AlphaFold2

The coarse-grained MD simulations were carried out using LAMMPS with the AWSEM model, under non-periodic shrink-wrapped boundary condition and Nose-Hoover thermostat control. The Hamiltonian of AWSEM contains both physical interaction terms and a bioinformatics-inspired memory term, *V*_*AWSEM*_ = *V*_*backbone*_ +*V*_*contact*_ +*V*_*burial*_ +*V*_*H−bond*_ +*V*_*FM*_ . Details of every term are described in Davtyan *et al*.^23^ and Papoian *et al*.^29^. AWSEM model has been applied to predict protein structures^23^, protein-protein interactions^24^, and in particular to histone dynamics such as chaperone-assisted histone dimer^30^, tetramer and octamer^14^, histone tails^31^, nucleosome^32^ and linker histone^33^.

To find low-energy conformations, simulated annealing was executed by decreasing the simulation temperature from 600 K to 200 K. The conformation that has the lowest energy is chosen as the prediction outcome. Ranging from 0 to 1, the order parameter *Q* (defined as 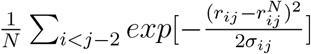 see S3.3 for details) was used to quantify the structural similarity to the native structure. To compute free energy profiles at a certain temperature, umbrella sampling was applied with *Q* as the reaction coordinate. WHAM^34^ was used to remove the potential bias. Protein sequences of all studied systems are provided in session S2. More AWSEM simulation details are in S3.1.

The all-atom MD simulations were performed using OpenMM 7.6.0^35^ on GPUs, with the Amber ff14SB protein model and the TIP3P water model. Initial conformations were either from the crystal structure, or predictions of AWSEM-MD and AF-Multimer. All the simulated systems were solvated in a 150 mM KCl solution. In total, eight molecular systems were separately simulated with two independent replicas for each. After standard energy minimization and equilibration, 800 ns were run for each replica, in total giving 12.8 microseconds of simulations. Results in the main context are from one replica. Complete all-atom results are in S3.2.

The AlphaFold-Multimer predictions for protein-protein complex were carried out using ColabFold^36^ on cloud service platform Google Colaboratory, with the fast homology search method MMseqs2^37^, the paired alignment and amber relaxation options.

### NMR and CD Experiments

Unlabeled and ^15^N labeled histones human H2A (type 2-A) and H2B (type 1-C) were expressed in *E. coli* and purified from inclusion bodies using cation exchange chromatography. Their correct mass was confirmed by mass spectrometry. All NMR experiments were performed at 23^*◦*^C on a 600 MHz Bruker Avance-III NMR spectrometer equipped with TCI cryoprobe. Proteins were dissolved at 100-200 *µ*M in 20 mM sodium phosphate buffer (pH 6.8) containing 7% D_2_O and 0.02% NaN_3_. NMR data were processed using TopSpin (Bruker Inc.)

Circular dichroism (CD) spectra of histone proteins at 0.40 mg/mL concentration in 20 mM sodium phosphate buffer at pH 6.8 were acquired on a Jasco J810 Spectro-Polarimeter using a Peltier-based temperature-controlled chamber at 25^*◦*^C and a scanning speed of 50 nm/min. A quartz cell (1.0 mm path length) was used. All measurements were performed in triplicate. To determine the secondary structure content, the CD data were analyzed using the DichroWeb server^38^. Two methods were used in parallel: (i) CDSSTR, a singular value decomposition (SVD)-based approach employing two datasets (7 and SMP180) from the DichroWeb server and (ii) K_2_D, a neural network-based algorithm trained using reference CD data^39^.

## RESULTS

### The Folding-upon-binding Mechnism of Histones

The individual folded monomers were not detectable in the past unfolding experiments of histone dimers or tetramers^10,11^. Our first aim was to investigate the molecular basis for this observation using AWSEM-MD. For that, we simulated 11 different histone monomers which include the eukaryotic canonical histones such as H2A and H2B, the variant histone CENP-A, and archaeal histones HmfA and HmfB. Histone tails were not included. Ten independent temperature annealing simulations were run for each monomer. Our simulation indicates that histone monomers are highly disordered at the tertiary structure level, as evidenced by the structural similarity measurement *Q* values less than 0.4 (Figure 2A, definition of *Q* see Methods), indicative of far-from-native conformations. The corresponding root-mean-square deviations (RMSD) from the native structure are more than 8 Å. Clustering analysis of the final conformations from different runs exhibits that no consensus tertiary structure was found (Figure S2). On the other hand, histone monomers exhibit stable secondary structure elements, especially the long *α*2 helix, which are similar to the native structure (Figure 2A). Without the stabilizing contacts from another monomer, the partially formed two short helices *α*1 and *α*3 of one histone are highly mobile, moving around the *α*2 helix, leading to significant tertiary disorder.

**Figure 2:**
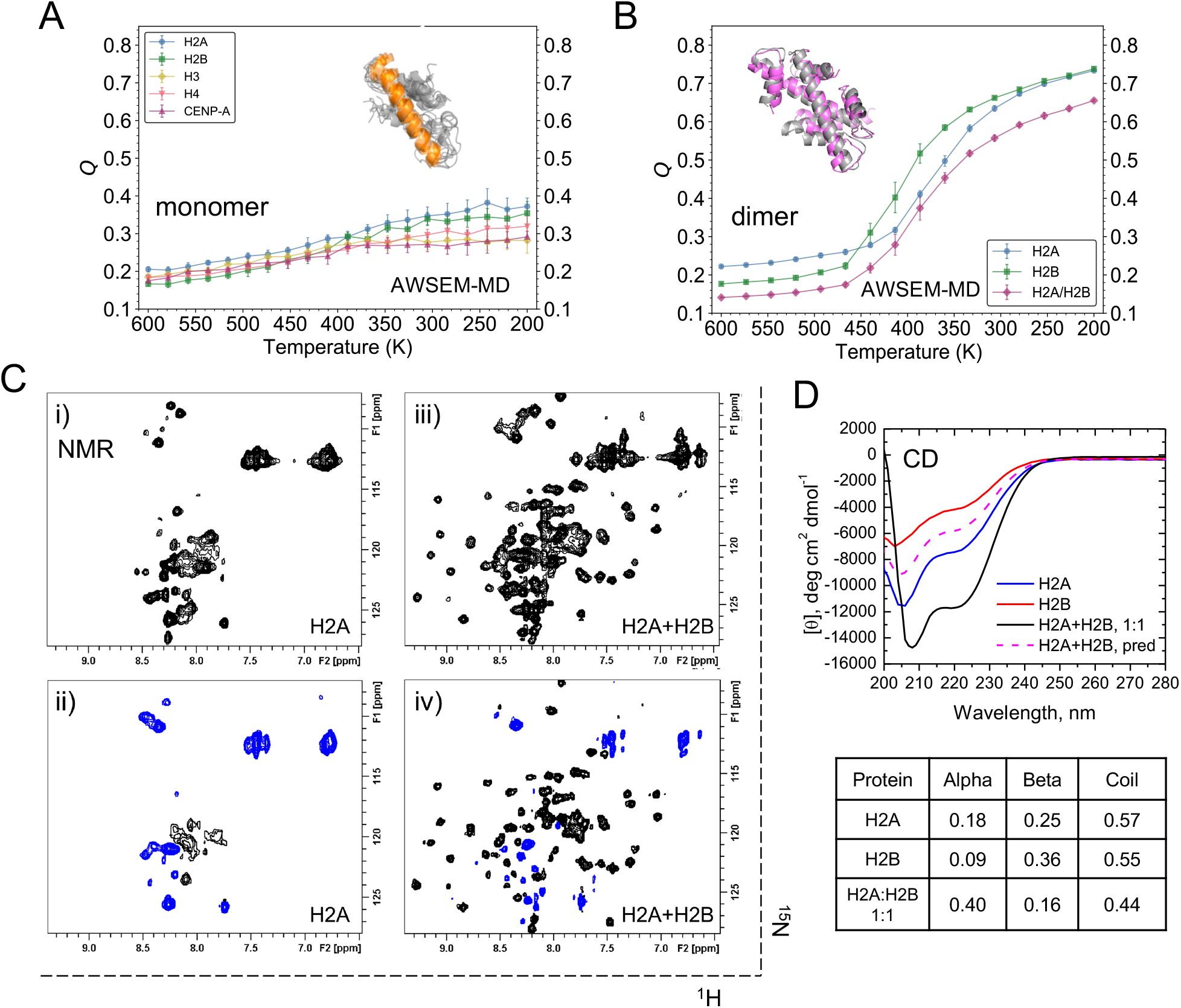
Histone heterodimers fold in MD simulations and NMR experiments, not monomers. (A) *Q* values as a function of the annealing temperature are plotted for simulated histone monomers, with the mean and standard deviation displayed as circles and error bars. Aligned final snapshots of H2A by *α*2 helix (orange) show that no stable tetiary structure formed. (B) *Q* analysis shows that the monomers of H2A and H2B (blue, green) cooperatively fold and bind into H2A/H2B dimer (magenta). The final conformation (magenta) is well aligned with its native state (gray). (C-D) NMR and CD studies of H2A and H2B upon their complex formation. (C) _1_H-_15_N NMR spectra of _15_N-labeled H2A alone (i) and in the presence of unlabeled H2B at a 1:1 molar ratio (iii). Heteronuclear steadystate _15_N{_1_H} NOE spectra with amide proton presaturation recorded for _15_N-labeled H2A alone (ii) and in the presence of unlabeled H2B at an equimolar ratio (iv). In these spectra, contours with positive intensities are colored black while negative intensities are blue. (D) CD spectra of H2A (blue) and H2B (red) alone and in an equimolar mixture (black). The total concentrations of the proteins are the same in all three cases. Also shown (dashed magenta) is the expected CD spectrum of the equimolar mixture of H2A and H2B if there were no structural changes in either protein. The table shows the secondary structure composition of the proteins estimated from these experimental CD data using K_2_D algorithm (see Methods for details).

We next applied a similar annealing protocol to investigate the folding of histones in the presence of their cognate binding partner. We calculated *Q* values of the entire dimer and of its component monomers. It shows that during the annealing run, *Q*_*dimer*_ increases roughly concurrently with *Q*_*monomer*_ (Figure 2B), indicating a clear structural transition wherein the two interacting monomeric chains cooperatively fold and bind. More than 95% native contacts were finally correctly predicted in the absence of any bias towards these contacts by the AWSEM force field (Figure S5). Analogous studies were undertaken for eukaryotic histones, histone variants and histone-like proteins including H2A/H2B, H3/H4, CENP-A/H4, archaeal histones (HMfA)_2_, (HMfB)_2_ and HMfA/HMfB, transcription factors dTAF_II_42/dTAF_II_62 and NF-YB/NF-YC (Figure S4, S5, S6, S7). All the results suggest that histones and histone-like transcription factors only fold upon binding with their partners. Interestingly, from the folding trajectories of the simulated histone dimers, it seems that histones first fold their long *α*2 helices, then form other *α* helices and higher-level inter-molecular tertiary structures (see the supplemental video).

To experimentally probe histone dimer folding, we carried out NMR and CD measurements on H2A and H2B. 1H-_15_N NMR spectrum of H2A alone (Figure 2C i) shows a narrow spread of NMR signals resulting in signal crowding in the region typical for amide signals of unstructured/unfolded proteins. The negative or close to zero signal intensities observed in the heteronuclear steady-state NOE spectrum of _15_N-labeled H2A recorded upon pre-saturation of amide protons (Figure 2C ii) are a clear indication that the protein is unstructured and highly flexible. Upon addition of unlabeled H2B, we observed a dramatic change in the _1_H-_15_N NMR spectra of _15_N-labeled H2A, wherein new signals (corresponding to the bound state) emerge and increase in intensity until they saturate at ca. equimolar H2B:H2A ratio (Figure 2C iii). Concomitantly, the unbound signals reduce in intensity and practically disappear at the saturation point. This behavior of the NMR signals, which exhibit essentially no gradual shifts, indicates that the binding is in slow exchange regime on the NMR chemical shift time scale. In contrast to the unbound state, the signals of _15_N-labeled H2A in complex with H2B (Figure 2C iv) show a significant spread, indicating that the bound state of H2A is well structured. Also, many H2A signals in the heteronuclear steady-state NOE spectra recorded at these conditions have positive intensities, which are characteristic of a well-folded state of the protein^40^. A similar behavior was observed for ^15^N-labeled H2B, which is unstructured in the unbound state and folds upon complex formation with H2A (Figure S9).

The NMR data presented above suggested that only with each other can H2A and H2B fold into a histone dimer with well-defined structures. To extend this analysis further, we performed CD measurements, which allow one to assess the helical content of a protein experimentally. The CD results demonstrate a significant increase in the helical content of these proteins upon formation of the H2A/H2B heterodimer (Figure 2D). Together, these experimental results indicate that in isolation H2A and H2B are disordered but adopt a well-defined tertiary structure upon binding to each other, which is consistent with a previous experimental study of H2A/H2B’s thermodynamic stability^10^. The 40% helical content of the human H2A/H2B heterodimer observed here is in excellent agreement with that for *Xenopus laevis* H2A/H2B^41^. Overall, our experiments and the above elaborated simulations are in qualitative agreement.

### FE Landscape Highlights a Stable Inverted Histone Dimer

Besides the native conformation, we observed an interesting non-native structure in the annealing simulations of core histone dimers. Compared to that of the native complex, the secondary structural elements of the non-native complex show very little differences but are arranged differently, leading to an inverted inter-molecular tertiary structure. The *α*1 helix of one histone is in proximity of the other’s *α*3 helix (Figure 3A, right), instead of two *α*1 helices close to each other as in the native structure (Figure 3A, left). If we define a direction pointing from N- to C-terminus for each histone, the angle between the two *α*2 helices in the inverted conformation is nearly complementary to that of the native. Not only in eukaryotic histones, this inverted conformation was also found in other simulated histone-fold systems, including archaeal histones and transcription factors (Figure S7). Interestingly, the native and inverted complexes are energetically comparable according to the AWSEM potential (Table S8).

**Figure 3:**
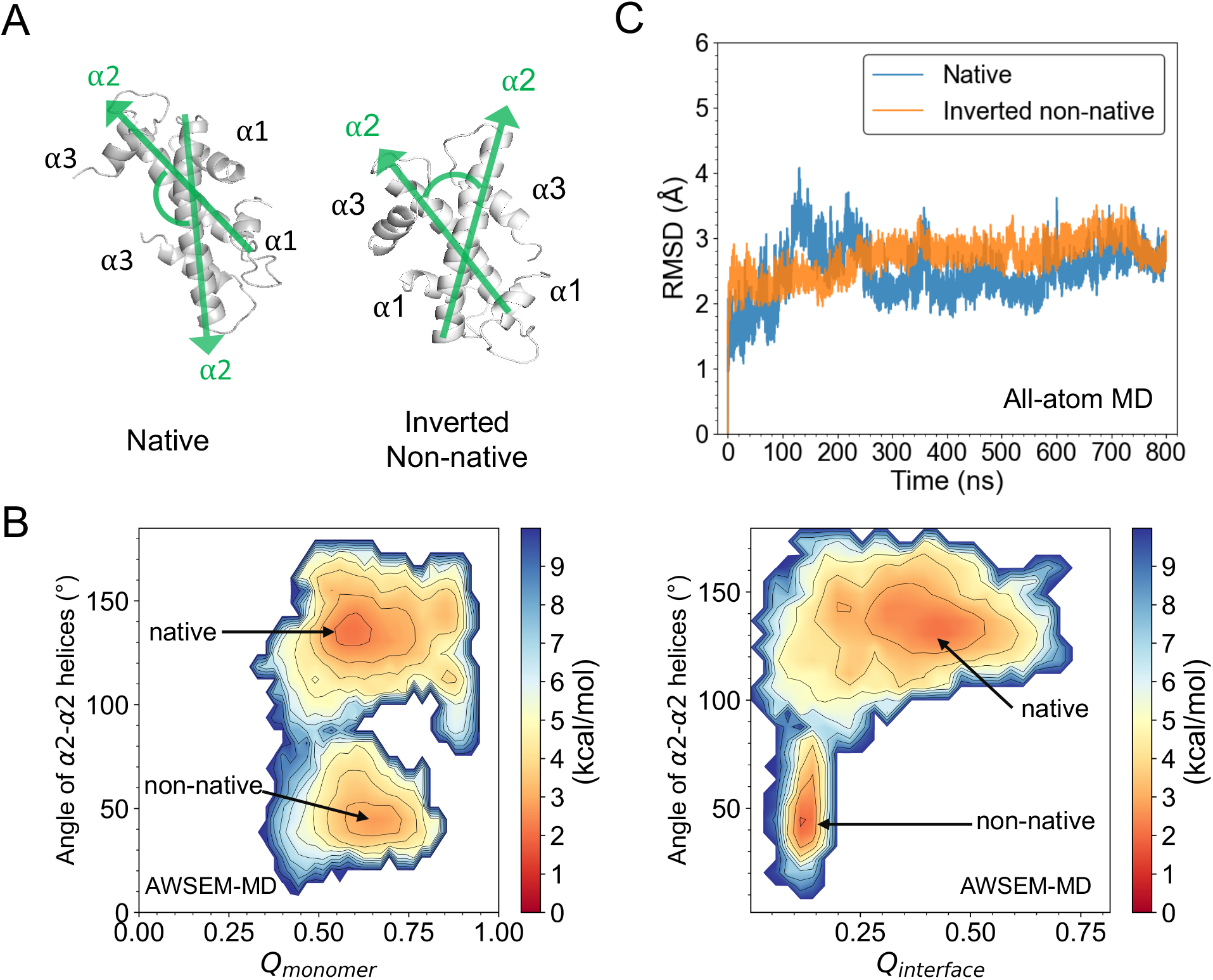
Inverted non-native conformation is found to be a stable formation of histone dimer. (A) The native and non-native conformation of H2A/H2B found in AWSEM simulations are shown. Their major difference is measured by the angle between the *α*2-*α*2 helices (green) with each vector arrow pointing from N- to C-terminal. (B) Free energy profiles of H2A/H2B are projected on *Q*_*monomer*_ and the angle of *α*2-*α*2 helices (left), and on *Q*_*interface*_ and the *α*2-*α*2 helices angle (right). Two energy minima are found, corresponding to the native-like and inverted non-native conformation respectively. (C) All-atom MD simulations show comparable stability of inverted non-native conformation to that of native structure of histone dimer.

To have a quantitative understanding of histone complex folding, we probed the FE landscape of H2A/H2B using umbrella sampling with AWSEM. The following three reaction coordinates were used to project the obtained FE profiles: the angle between two *α*2 helices to describe the general inter-chain geometry (Figure 3A), *Q*_*monomer*_ to show how well histone monomer is folded, and *Q*_*interface*_ to show the nativeness of formed histone dimer interface. In both of the FE landscapes (Figure 3B), there are two major basins. The first basin (top one in both panels) is located at the *α*2-*α*2 angle of 140^*◦*^, both *Q*_*monomer*_ and *Q*_*dimer*_ high, all of which are consistent with the native conformation. The second basin (lower one in the figure) has the *α*2-*α*2 angle of 40^*◦*^, high *Q*_*monomer*_ but low *Q*_*dimer*_, indicating the formation of native-like monomers and a “mismatched” binding interface. The representative structures at the two basins show that they are the native-like and the inverted non-native conformations as we found in previous annealing runs. The native basin seems to be significantly broader, suggesting entropic stabilization. The free energy barrier between the two states is relatively high at ∼8-9 kcal/mol.

To further test the structural stability of the newly-found non-native histone dimer, we took the final snapshot of its inverted conformation from AWSEM prediction and used that as the initial conformation to perform extensive all-atom MD simulations in explicit solvent (see Methods and S3.2 for details). The subsequent RMSD analysis indicates that the non-native structure reached a steady state and maintained remarkable structural stability (Figure 3C), which is comparable to the simulation started from the native struture. More results are in session S6.

### Sequence Symmetry Explains the Inverted Conformation

Next, we explored the molecular interactions that potentially promote the non-native histone dimeric complex formation. It is known that in the native complex, *α*1, *α*2, and *α*3 helices of one histone interact with the other histone through hydrophobic interactions (Figure 4A). These hydrophobic residues are typically conserved among all histone structures (Figure S2). Interestingly, we found that in the inverted non-native complex these interactions are still favored when swapping *α*1 and *α*3 helices. To further explain its biological basis, we reversed the histone sequences from C- to N-terminus, and carried out sequence alignments between the reversed and normal order histone sequences (Figure 4B). Consequently, the multiple sequence alignments reveal a symmetric distribution of hydrophobic residues among histones and their head-tail reversed sequences. Thus, one may expect that in an N-C-terminal inverted arrangement, histones should still be able to maintain the required hydrophobic interactions for a stable formation due to the conserved symmetry of hydrophobic positions.

The finding of head-tail sequence symmetry and inverted histone dimer well support the previously proposed histone evolution hypotheses. The latter posits that histones may have arisen through the duplication and differentiation of a primordial helix-strand-helix motif ^7,42^ (Figure 4C i-ii), and domain swapping may have triggered the subsequent dimerization of two histone monomers^26,43^ (Figure 4C iii). Thus, it is possible that the conservation of hydrophobic positions found in histone sequences, no matter in a normal or reversed order, originated from the same ancestral peptide. Meanwhile, the conserved hydrophobicity in the four segments of two histones explains the rationale of two alternative ways of the crossing forming the hand-shake motif for the characteristic of the histone fold.

**Figure 4:**
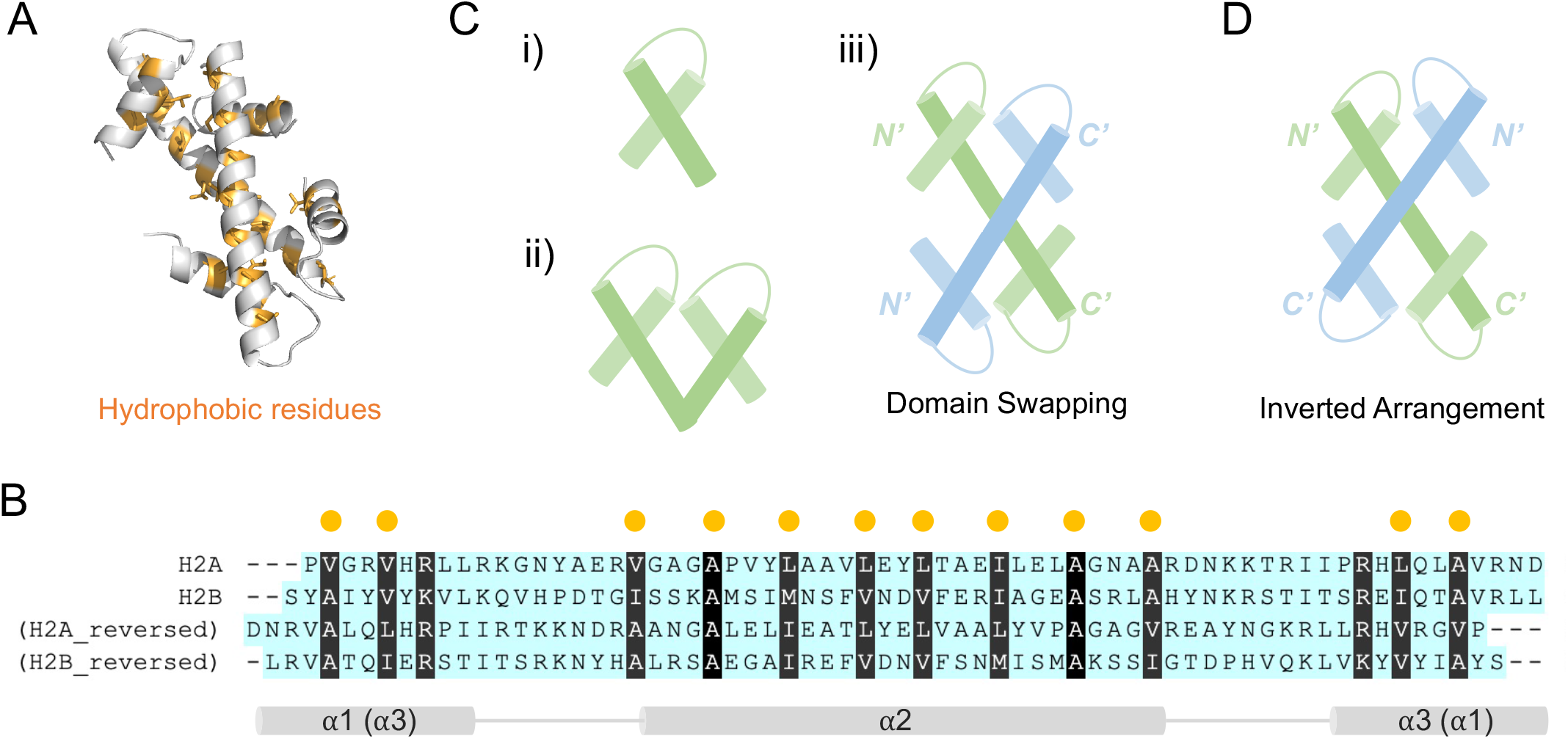
Sequence symmetry of hydrophobic residues explains the predictability of inverted histone-fold structure. (A) Hydrophobic interactions (orange) dominate the formation of binding interface of H2A/H2B (silver). (B) Protein sequence alignments of histones H2A, H2B and their reversed sequences highlight the symmetrical distribution of hydrophobic residues. Hydrophobic residues are particulary marked. (C) Cartoon schemes illustrate previously proposed histone evolution hypotheses: i) histones may originate from one single helix-strand-helix structural motif; ii) duplication, differentiation and fusion of two helix-strand-helix peptides result in one protein; iii) domain swapping between two proteins (colored in green and blue) forms a histone-fold structure. (D) The inverted arrangement based on hydrophobic interactions could be an alternative formation of histone-fold structure.

In summary, our finding of energetically favorable native and inverted dimers points to the hidden ancestral symmetry of histone protein sequences. Analogous sequence symmetry in an unrelated protein, the Rop dimer, was previously suggested to result in a double-funnel free energy landscape^44,45^. In this context, our results suggest that histone chaperone, or other molecular factors and regulators may be needed to facilitate correct folding of histone dimers, which, in turn, would ensure that the correct binding interface is exposed for their subsequent interactions with DNA and other histone proteins.

### AWSEM-MD and AlphaFold-Multimer Predict Eukaryotic Homodimer Structure

The above observed inverted non-native histone-fold conformation was also observed in archaeal histones (HmfA)_2_, (HmfB)_2_, HmfA/HmfAB, and transcription factors dTAF_II_42/dTAF_II_62 dimer and NF-YB/NF-YC dimer (Figure S6). These results indicate a well-conserved folding mechnism of histone-fold structures regardless of species and function distinctions, which prompted us to investigate the hypothetical eukaryotic homodimeric histones. As known, a significant difference of histone oligomers between Eukarya and Archaea is that eukaryotic histones naturally exist as heterodimer while in Archaea both homodimer and heterodimer histones are prevalent. In early biochemical investigations of histone-histone interactions, sedimentation experiments showed that eukaryotic homotypic histones may exist at high histone and salt concentrations^46,47^, however, without additional structural elaboration.

Based on the above findings, we used AWSEM annealing simulations to predict the structure of a putative H2A/H2A homodimer (histone core only, N- and C-termini were not included). Among the predicted structures, we found both native-like and the above mentioned inverted conformations. The RMSD between native-oriented H2A/H2A complex and the native H2A/H2B structure is 6.0 Å. We then applied AlphaFold-Multimer^27,28^ in ColabFold^36^ to carry out analogous complex structure prediction for H2A/H2A. Interestingly, both native-oriented and inverted structures were predicted as well, consistent with our predictions from the AWSEM model. The top-scored prediction from AF2 is a native-like histone-fold structure, with a RMSD of 2.6 Å to the native heterodimer structure of H2A/H2B. The structural alignments to the H2A/H2B native structure (Figure 5A) indicate that predictions from two totally different methodologies, AWSEM and AF2, agree well. We note that we have used two models of AF2-based prediction tools. Details of the two versions’ performance and related discussion about protein complex prediction are in session S7.

**Figure 5:**
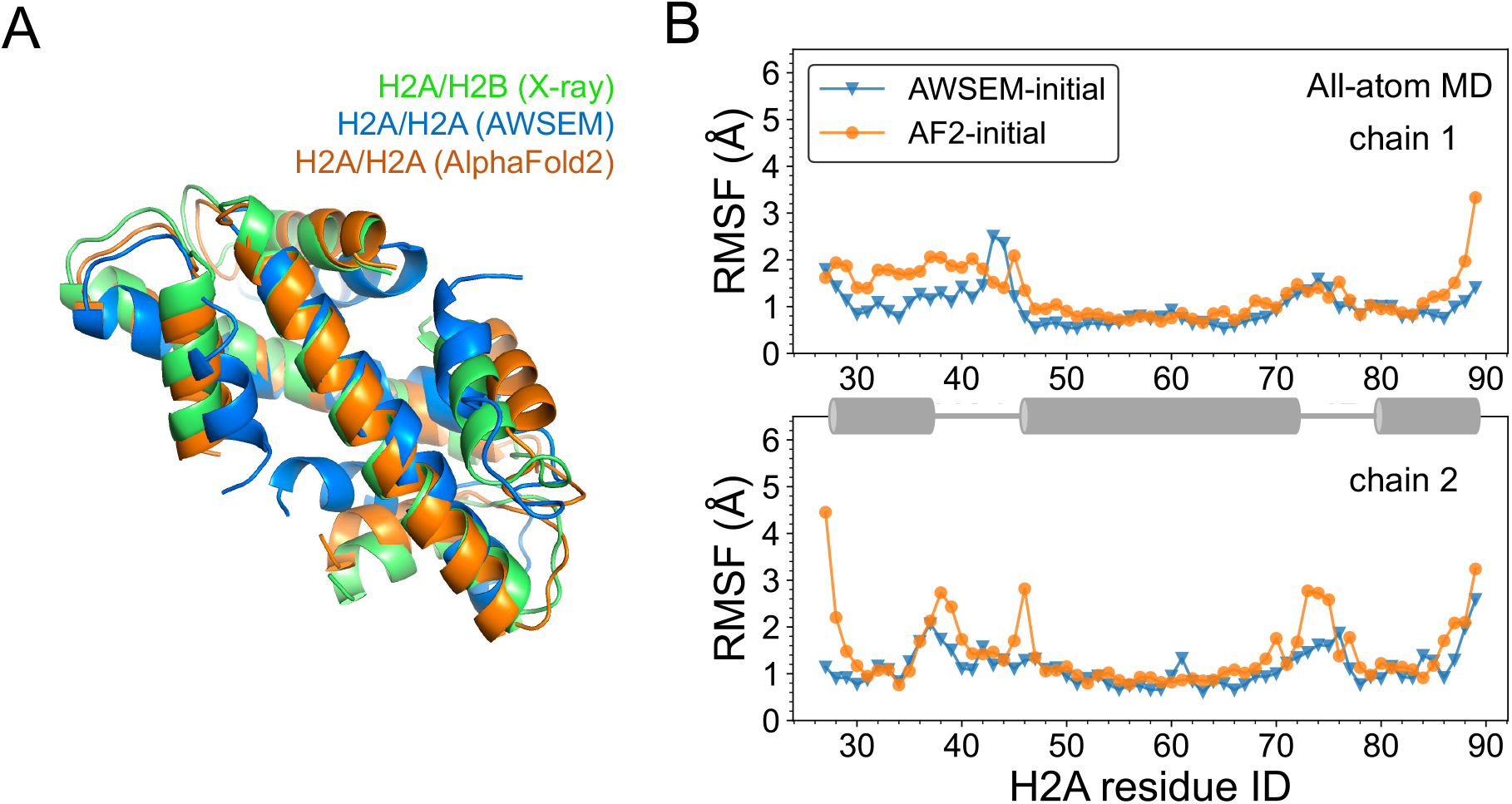
Predicted structures of eukaryotic histone homodimer. (A) AWSEM and AlphaFold2 predicted homodimer structures of H2A/H2A align well with the native heterodimer structure of H2A/H2B (colored in blue, orange, and green). (B) RMSF analysis of all-atom simulations demonstrates comparable stabilities of AWSEM- and AlphaFold2-predicted homo-complex structures (in blue and orange). RMSF of two chains are plotted separately and helix regions are animated by cartoon in grey.

To examine the structural stability of the H2A/H2A histone homodimer, we further carried out explicit solvent all-atom simulations starting with the two initial conformations predicted by AWSEM, and AF2, respectively. We structurally aligned the simulation trajectory snapshots to the initial conformation, and calculated the root-mean-square-fluctuation (RMSF) of the C_*α*_ atoms to quantify local residual fluctuations (Figure 5B). The RMSF analysis displays comparable fluctuations between the two simulations on average within 3 Å. The RMSF analysis also shows that the loop regions of histone homodimer are in general more flexible than helical regions, as we expected.

### The Role of Histone Tails in Regulating Homodimer Formation

The above results suggest that eukaryotic histone homodimers may be sufficiently stable in the absence of histone tails. This finding points to the possibility of histone tails tilting the balance between the homo- and heterodimer towards the latter. Hence, we next predicted the full-sequence histone homodimers using simulated annealing in AWSEM, and subsequently evaluated their structural similarities to the native heterodimer H2A/H2B (only for the histone fold core part) via the structural similarity measure, *Q*_*dimer*_ (Figure 6A). From the best five predicted structures for each group, we can see that full-sequence H2A/H2A has significantly lower *Q*_*dimer*_ than H2A/H2A without histone tails (referred to “truncated H2A/H2A” hereafter), indicating a disrupting role of histone tails in the formation of the histone homodimer. This inhibitory effect of histone tails was not observed in the formation of H2A/H2B complex. Interestingly, instead, the full-length H2A/H2B shows slightly better *Q*_*dimer*_ values than what were found for truncated H2A/H2B. This result is consistent with a previous experimental study showing that H2A/H2B N-terminal tails may speed up the correct folding of H2A/H2B^48^.

**Figure 6:**
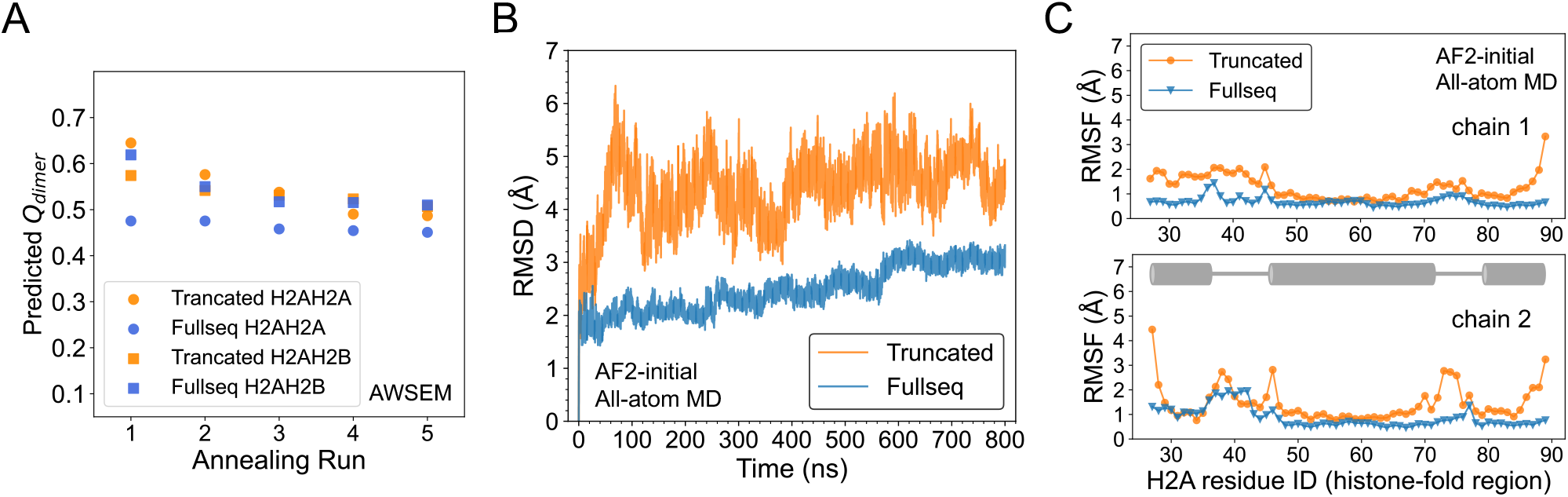
Histone tails disfavor the formation of histone homodimer, yet stabilize the histone fold once formed. (A) AWSEM-predicted trancated and full-sequence homodimer H2A/H2A are assessed by *Q*_*dimer*_ of the histone-fold region (circle in orange and blue) and compared with predictions of trancated and full-sequence heterodimer H2A/H2B as a control (square in orange and blue). (B-C) All-atom simulations of AlphaFold2-predicted truncated (orange) and full-sequence (blue) H2A/H2A homodimer are analyzed through root-mean-square-deviation (RMSD) (B) and root-mean-square-fluctuation (RMSF) (C) for the histone-fold region. In (C), the two chains of H2A/H2A are plotted separately and their helix regions are animated by a cartoon diagram.

Moreover, we used AlphaFold-Multimer to predict the full-sequence structure of H2A/H2A. The obtained structure is very similar to that of H2A/H2B, with a *Q*_*dimer*_ of 0.7. We then performed all-atom MD simulations using the predictions of AF-Multimer for H2A/H2A homodimer with and without histone tail (truncated) as initial conformations. Interestingly, the full-sequence H2A/H2A has lower RMSD on average compared to its initial conformation than truncated H2A/H2A (Figure 6B). RMSF analysis (Figure 6C) shows that the flexibility difference between the two systems mainly appears at the terminal residues of the histone-fold core and the loop regions. In summary, once the H2A/H2A has formed a histone-fold core, the disordered tails may potentially stabilize the histone-fold by forming interactions between tails and histone-fold core.

## DISCUSSION

From a biology perspective, our study shows that the eukaryotic histone dimers not only have the same structural motif as their ancestor proteins, but also the similiar folding mechanism. However, an interesting consequence of eukaryotic histones having forked from the archaeal tree is that eukaryotic histones have tails and do not form homodimers. The length of histone tails and their post-translational modifications vary quite widely among variants found in eukaryotic organisms, playing important roles in determining the linker DNA and nucleosome interaction. ^33,49^ In this work we suggest that the addition of histone tails impedes the formation of eukaryotic homotypic histones. In addition, histones’ evolution not only expanded the structural variability but also dramatically enhanced the functional repertoire of histone dimers, for example in DNA repair^50^, enhancer activity^51^, and centromere function^52,53^. Possible new functions may arise from distinct post-translational modifications on each monomer^54–56^.

Our current finding of relatively stable inverted dimers suggests that chaperones, either protein, DNA or RNA, may be generally needed to assist proper histone folding. The result that both archaeal and eukaryotic histones cannot fold into a stable monomeric structure also indicates the general necessity of chaperones or other binding partners to histone monomers. Indeed, a number of eukaryotic histone chaperones have been found in recent decades^57–62^, such as Asf1 and MCM2 for H3/H4, Nap1 and MBD for H2A/H2B. Yet, most chaperones reported so far bind with dimeric histone complex. A recent exeperimental work by Pardal and Bowman reports that histone H3 and H4 can be separately chaperoned and transported by Imp5-H3 and Imp5-H4, and then get dimerized with other chaperones such as NASP and HAT1-RBBP7^63^. This work suggests that the individual histone proteins H3 and H4 may be structurally stablized by Imp5. Besides, the revealed complex regulation network of histone chaperones can be a consequence of the typically existent conservation and symmetry that underly all histones and their binding partners both in sequence and structure.

Interestingly, no archaeal histone chaperone has yet been reported. Thus, we anticipate the existence of archaeal histone chaperones or new chaperoning function from known proteins. However, we do not exclude the possibility that extreme ionic or temperature conditions which define the extremophile habitat, might somehow prevent the non-native association of archaeal histones. Furthermore, the ratio of native to non-native complexes varied among the different dimers that we simulated (Figure S7). This implies that the sequence and structural diversities of histones, the varying lengths of histone tails, and post-translational modifications may all affect their overall folding dynamics. For instance, previously we reported that histone monomers H3 and H4 play distinct structural and functional roles in their heterodimer H3/H4 complex^30^. It is possible that a chaperone would interact with only one histone monomer and primarily assist its folding. HJURP mainly interacting with CENP-A in the CENP-A/H4 dimer represents one such salient example. Consequently, one may anticipate close co-evolution between histone monomers and chaperones.

## CONCLUSIONS

In this work, we used computational approaches supplemented with NMR and CD experiments to investigate the folding mechanism of histones. We found that a cooperative folding-upon-binding principle widely applies to canonical and variant eukaryotic histones, archaeal histones as well as histone-like transcription factors. Moreover, we report that histone cores without tails form a non-native dimeric complex, in which on the tertiary structure level, an inverted arrangement is energetically nearly competitive with the native conformation. Further sequence analysis indicates that this surprising non-naive stable formation could be a consequence of the ancient sequence symmetry underlying all histone proteins. Finally, we used AWSEM, AlphaFold-Multimer and all-atom MD simulations to predict possible formations of eukaryotic homodimers. Our results show that histone tails play an important role in regulating the formation of eukaryotic homodimer or heterodimer. Without tails, eukaryotic histones form stable homodimers in both AWSEM and AlphaFold-Multimer predictions, while with tails AWSEM simulations indicate an impeded formation of histone homodimers. Overall, we molecular details of histone folding dynamics, providing a more comprehension in guarding histone homeostasis.

## Acknowledgement

The authors thank Drs. David Winogradoff, Guang Shi and Daniël Melters for helpful feedbacks, Carlos Castañeda for help with initial NMR measurements, and Tingting Yao for kindly providing plasmids for histone expression. H.Z. also thanks to the NCI-UMD partnership program for Integrative Cancer Research. This work was supported by the Ann G. Wylie Dissertation Fellowship (H.Z.), the Amazon Web Services Artificial Intelligence Award (G.P.), the NIH Intramural Research Program (Y.D.), and the NIH grant GM065334 (D.F.).

## Supporting Information Available

The Supporting Information contains the following contents:.

- Complete information for all simulated protein structures (S1, S2)
- Method details of AWSEM and all-atom MD simulations (S3)
- Supplemental AWSEM results of all simulated structures (S4, Fig. S2-S7, Table S8)
- More NMR experiment results (Fig. S9)
- All-atom simulation results from another replica (S6, Fig. S10-S12)
- Structures of H2A/H2A, H2B/H2B predicted by AlphaFold-Multimer (Fig. S13-S15)
- Discussion about protein complex structure prediction (S7)

## Simultation Movies Available

Folding trajectories from AWSEM-MD simulations are provided respectively for the native and inverted non-native conformation of H2A/H2B (eukaryotic histones), and for the native and inverted non-native conformation of HmfA/HmfA (archaeal histones). Together, these videos highlight the symetrey and asymetry behind histone folding across the eukaryotes and archaea.

