## Supplemental Information for "The Symmetry and Asymmetry Behind Histone Folding Across Eukarya and Archaea"

### Contents

**S1 Polymer Scaling Analysis** **S3**

**S2 Sequences of Studied Proteins** **S4**

**S3 Method and Analysis Details** **S5**

S3.1 AWSEM-MD Simulations . . . . . . . . . . . . . . . . . . . . . . . . . . . . S5

S3.2 All-atom MD Simulations . . . . . . . . . . . . . . . . . . . . . . . . . . . . S6

S3.3 Order Parameter $Q$ . . . . . . . . . . . . . . . . . . . . . . . . . . . . . . . S7

**S4 Supplemental AWSEM Simulation Results** **S9**

**S5 Supplemental NMR Results** **S16**

**S6 Supplemental All-atom Simulation Results** **S17**

**S7 Discussion for Histone Folding/Binding Mechanism and Complex Structure Prediction** **S19**

**References** **S22**

### S1 Polymer Scaling Analysis

Our computer simulations and NMR and CD experiments suggest that histone dimers should be considered as an independent folding unit. From a perspective of polymer physics, the radius of gyration for a polymer chain ($R_g$) approximately follows the scaling relation: $R_g \sim \alpha N^\nu$, where $N$ is the number of bond segments (*i.e.* the degree of polymerization) of the chain, $\alpha$ is the linear slope, and $\nu$ is the scaling exponent. Dima and Thirumalai[1] estimated this scaling relation for proteins after analyzing a large dataset of monomeric protein structures. Based on their obtained values of $\alpha \simeq 3$ and $\mu \simeq 1/3$, we fitted the $R_g$ and $N$ of the X-ray crystal structures of histone monomers and dimers (Figure 1). We found that all the histone monomers have higher $R_g$ than expected for a globular protein with the same residue length $N$. Histone dimers, on the other hand, follow very well the $R_g$ trend of single domain proteins, again supporting the idea that histone dimers represent a single folding unit.

### S2 Sequences of Studied Proteins

```
     H2A -------PVGRVHRLLRKGN---YAERVGAGAPVYLAAVLEYLTAEILELAGNAARDNKKTRIIPRHLQLAVRND------------------ 65
     H2B ------SYAIYVYKVLKQVH---PDTGISSKAMSIMNSFVNDVFERIAGEASRLAHYNKRSTITSREIQTAVRLLLPGELAKHAVSE------ 78
      H3 ELLIRKLPFQRLVREIAQDFK--TDLRFQSSAVMALQEASEAYLVALFEDTNLCAIHAKRVTIMPKDIQLARRIRGERA-------------- 77
      H4 RDNIQGITKPAIRRLARRG----GVKRISGLIYEETRGVLKVFLENVIRDAVTYTEHAKRKTVTAMDVVYALKRQGRT--------------- 74
   CENPA HLLIRKLPFSRLAREICVKFTRGVDFNWQAQALLALQEAAEAFLVHLFEDAYLLTLHAGRVTLFPKDVQLARRIRGLEEGLG----------- 82
    Hmfa ----GELPIAPIGRIIKNA----GAERVSDDARIALAKVLEEMGEEIASEAVKLAKHAGRKTIKAEDIELARKMFK----------------- 68
    Hmfb ----MELPIAPIGRIIKDA----GAERVSDDARITLAKILEEMGRDIASEAIKLARHAGRKTIKAEDIELAVRRFK----------------- 68
dTAFII42 ---------PKDAQVIMSILKELNVQEYEPRVVNQLLEFTFRYVTSILDDAKVYANHARKKTIDLDDVRLATEVTLD---------------- 68
dTAFII62 MLYGSSISAESMKVIAESI----GVGSLSDDAAKELAEDVSIKLKRIVQDAAKFMNHAKRQKLSVRDIDMSLKV------------------- 70
   NFY-B ----IYLPIANVARIMKNAIP--QTGKIAKDAKECVQECVSEFISFITSEASERCHQEKRKTINGEDILFAMSTLGFDSYVEPLKLYLQKFRE 87
   NFY-C ------LPLARIKKIMKLDE---DVKMISAEAPVLFAKAAQIFITELTLRAWIHTEDNKRRTLQRNDIAMAITKFDQFDFLIDIVPR------ 78
```

Figure S1: **Multiple sequence alignments (MSA) for all simulated proteins.** MSA of the histone and histone-like proteins in this study show that the conserved residues (highlighted in black shade) are mostly hydrophobic and located near the dimer interfaces.

### S3 Method and Analysis Details

#### S3.1 AWSEM-MD Simulations

In this work, we used the AWSEM[2] model to simulate all the histone and histone fold protein (HFP) systems. The parameters in AWSEM were tuned such that the simulated melting temperature (*i.e.* the temperature at which both the folded and unfolded states are equally populated at equilibrium) of histone dimers is around 350 K, as observed in experiments. In addition, we employed an AWSEM-featured bioinformatic term called "fragment memory", using available protein segments as local structural bias. In histone/HFP monomer annealing simulations, the biasing segments were selected from proteins in the PDB which share similar local amino acid sequences to the histone monomers, equivalent to the "homologue allowed" structural library used in Davtyan *et al.*[2] In the dimer simulations, the local memory fragments were selected from the X-ray crystal structures, which still only provide local structure information, but not tertiary contacts within each monomer and between monomers (as used in previous protein binding studies with AWSEM[3]). The length of a fragment is typically from 3 to 9 residues. To maintain a reasonable contact region, a weak distance constraint in the format of harmonic potential was applied between the centers of mass of two simulated monomers (the spring constant $k = 0.02$ kcal/mol/Å$^2$).

We ran AWSEM simulations using the open-source molecular dynamics software, LAMMPS[4] (version 9Oct12), with non-periodic shrink-wrapped boundary condition and the Nose-Hoover thermostat. The simulation time step was set as 5 femtoseconds. All annealing simulations started from the completely unfolded state, and then were slowly cooled down from 600 K to 200 K. The simulation time of a production run is $1 \times 10^7$ steps. Ten independent runs with different initial states and velocities were performed for each system. The native conformations were taken from the corresponding X-ray crystal structures (PDB: 1AOI[5] for histone H2A/H2B and H3/H4; PDB: 3R45[6] for CENP-A/H4; PDB: 1B67 and 1A7W[7] for archaeal histone HmfA and HmfB; PDB: 1TAF[8] for dTAF$_{\mathrm{II}}$; PDB: 1N1J[9] for

NF-Y). In current study, we focus on understanding the histone fold, so histone tails and N- and C-terminal helices are typically excluded in our simulations. The sequences of proteins used here can be found in the following multiple sequence alignment figure (Figure S1).

To calculate the free energy profile, we used umbrella sampling. The $Q$ of the simulated molecules is chosen as the reaction coordinate. We set up 19 umbrella windows along $Q$ ranging from 0 to 1. A harmonic potential around each $Q_0$ was added to the total Hamiltonian as in the equation $V(Q) = \frac{\kappa}{2}(Q - Q_0)^2$. The spring constant $\kappa$ that we used here is 1000 kcal/mol/Å$^2$. In each window, the initial conformation is prepared by annealing under the $Q_o$ potential biase procedures from 450 K to 250 K. The final umbrella sampling is at 300 K. Weighted histogram analysis method (WHAM) was used to remove the potential biase and calculate the free energy profiles.

All used protein sequences in this study are downloaded from UniProtKB database[10] and aligned using MUSCLE v3.8.31[11]. The generated multiple sequence alignments are visualized by an online tool, Multiple Align Show, $https://www.bioinformatics.org/sms/multi\_align.html$. Structure and simulation conformation figures used in this work are generated by PyMOL Molecular Graphics System Version 2.3.2, Schrödinger LLC.

#### S3.2 All-atom MD Simulations

The all-atom simulations were performed in the high-performance MD engine OpenMM 7.6.0[12], with the input files prepared by CHARMM-GUI[13]. The atomic force fields used in this work include Amber ff14SB force field for protein, the TIP3P water model for solvent, and the Joung/Cheatham ion parameters[14] for TIP3P water. Particle-Mesh-Ewald (PME) electrostatics and switched Leonard-Jones interactions with a cutoff distance of 12 Å were used in all the simulations. The protein systems to be simulated were solvated in 150 mM KCl solution, with periodical boundary conditions of a minimum distance of 12 Å. The timestep was set up at 2 fs. The energy minimization takes 50,000 timesteps, followed by equilibration under the NVT ensemble at 300 K for 500,000 timesteps. Hereafter the production runs were performed for 400,000,000 steps (800 ns) under the NPT ensemble, using Langevin thermostat and MonteCarlo barostat. Along each trajectory, atomic coordinates were saved every 25,000 steps.

For each simulated system, we run the abovementioned process for two independent replicas using different random seeds for their initial velocity states. The simulated systems are: 1). H2AH2B from the X-ray crystal structure (PDB: 1AOI) (trancated-sequence, chain A: P26-D90, chain B: Y34-L98); 2). AWSEM-predicted H2AH2B inverted non-native structure (trancated-sequence, chain A: P26-D90, chain B: Y34-L98); 3). AWSEM-predicted H2AH2A homodimer structure (trancated-sequence, P26-D90, P26-D90); 4). AlphaFold2-predicted H2AH2A homodimer structure (trancated-sequence, P26-D90, P26-D90); 5). AlphaFold2-predicted H2AH2A full-sequence homodimer structure (G4-K11, G4-K11); 6). Full-sequence H2AH2B from X-ray crystal structure (PDB: 1AOI) (G4-K118, K24-K122). In total, we run 9600 ns all-atom explicit-solvent simulations for different histone dimer structures.

Before running all-atom simulations, conformations from AWSEM predictions were refined using FoldX (v5.0)[15] to repair and optimize the side-chain atomic structures, and then using Chiron[16] to further remove steric clashes. Conformations from AlphaFold2 predictions were relaxed with amber force fields as implemented in the AlphaFold2 protocal.

#### S3.3 Order Parameter $Q$

To quantitatively describe the similarity between simulated and native structures, we used the order parameter $Q$ defined as in Davtyan *et al.*:[2]

$$Q = \frac{1}{N_{pairs}} \sum_{i<j-2} \exp\left[-\frac{(r_{ij} - r_{ij}^{N})^2}{2\sigma_{ij}}\right] \tag{S1}$$

where $N_{pairs}$ is the number of pairs in the summation, $r_{ij}$ is the instantaneous distance between $C_\alpha$ atoms of residues $i$ and $j$, $r_{ij}^N$ is the same distance in the native structure, and $\sigma_{ij} = (1 + |i - j|)^{0.15}$ represents the resolution of distance difference.

The range of $Q$ is from 0 to 1. A higher $Q$ value means that the simulated conformation is more similar to the native structure. Note that the group of atoms included for computing $Q$ can be customized. In the main text, we computed $Q_{monomer}$ using the $C_\alpha$ atoms only within a monomer, while $Q_{dimer}$ was calculated using the entire dimer.

### S4 Supplemental AWSEM Simulation Results

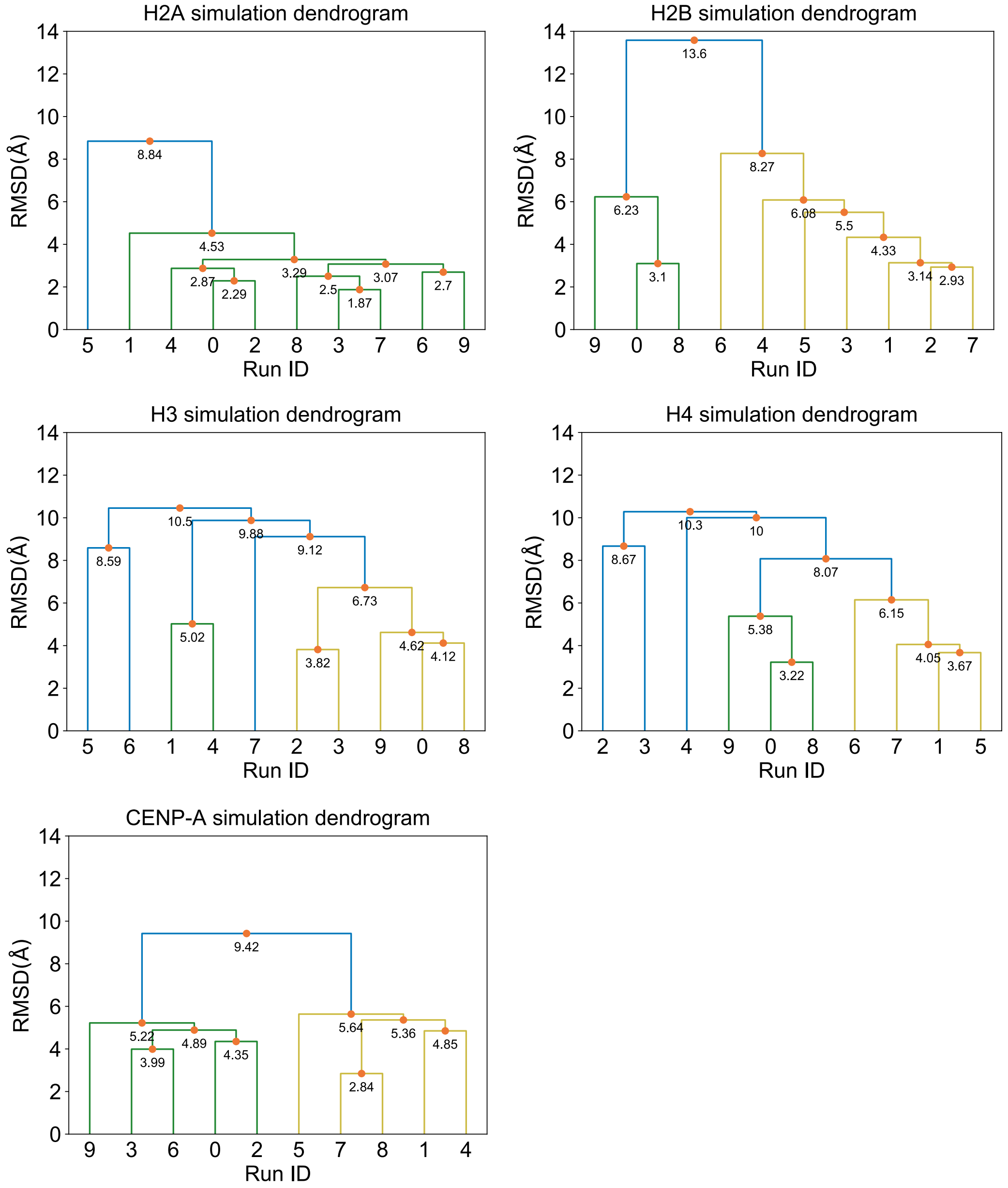


Figure S2: **Clustering analyses for the final snapshots of histone monomer simulations.** Shown here are the clustering analysis results based on the pairwise RMSD among ten independent annealing runs for each histone monomer. No consensus structure was found.

We implemented analogous AWSEM simulations for other histone monomers including archaeal histone HMfA and HMfB, histone-like transcription factor monomer dTAF$_{II}$42 and dTAF$_{II}$62, NF-YB and NF-YC, and their histone pairs including eukaryotic histones

H3/H4, histone variant CENP-A/H4, archaeal histones $(HMfA)_2$, $(HMfB)_2$ and heterodimer HMfA/HMfB, transcription factors dTAF$_{II}$42/dTAF$_{II}$62 dimer and NF-YB/NF-YC dimer. We confirmed similar observations that monomeric histones (or histone-like protein) cannot be folded (Figure S3). All simulated complexes— histones and their binding partners—have similar folding-upon-binding patterns for all (Figure S5, S6) and an inverted non-native formation besides their native ones (Figure S7).

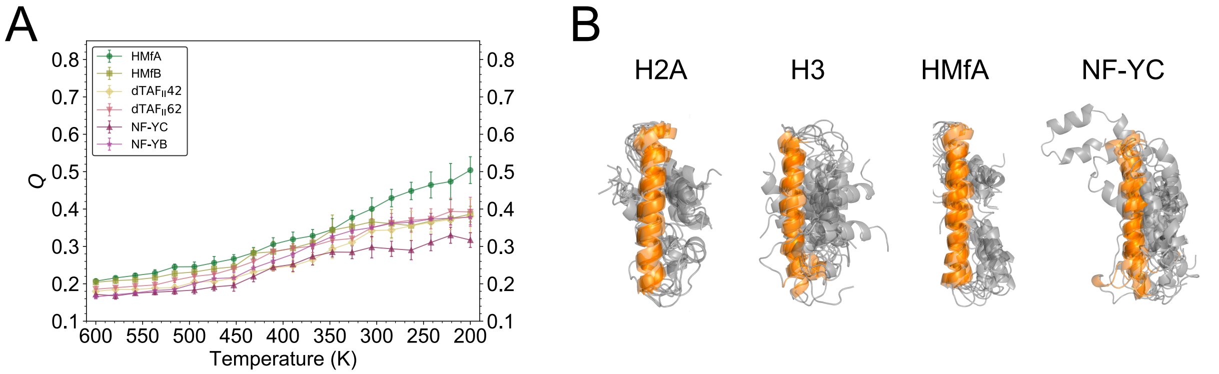


Figure S3: **Simulations of archaeal histone monomer and histone-like protein monomer** (A) Complementary to Figure 1B in the main context, here are simulation results for monomers of archaeal histone and two types of transcription factor dTAF and NF-Y. (B) Structure alignments based on the longest $\alpha 2$ helix (orange) for monomer simulation of H2A, H3, HMfA and NF-YC. Only the final conformations in corresponding annealing runs are included.

It is noteworthy that previous studies used biochemical measurements to demonstrate the formation of archaeal heterodimers HMfA/HMfB *in vitro*[7], which, however, was not structurally characterized. Here, based on our above-mentioned findings (Figure 2 and S5, S6), we hypothesized that archaeal histone heterodimers adopt a structure and binding-folding mechanism analogous to other histone dimers and the archaeal histone homodimer. To test this idea, we took HMfA and HMfB monomer structures from their homodimer complexes (PDB: 1B67 and 1A7W)[7], and combined them into a heterodimer, which was subsequently studied using simulated annealing. These computational experiments indicate that the heterodimer indeed also follows the same structural and folding patterns as other histones (Figure S4). We further applied umbrella sampling and calculated its FE profiles. It turns that similar two states with complimentarily orientated are found in two energy basins (Figure S4.B). The energy barrier between the two minimum states is similarly 9 Kcal/mol.

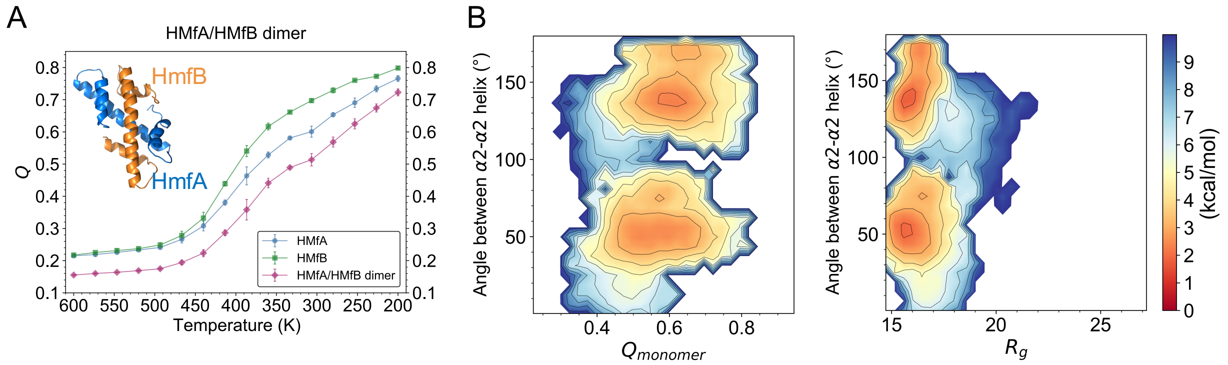


Figure S4: **Predicted heterodimer of archaeal histones follows the same folding mechanism.** (A) $Q$ value of the archaeal histone HMfA (blue) and HMfB (green), and its dimer (magenta), and our computational prediction of archaeal heterodimer HMfA/HMfB structure are shown. (B) Free energy profile calculated at 300 K are projected on $Q_{monomer}$ and the $\alpha$2-$\alpha$2 angle (left), and radius of gyration $R_g$ and the $\alpha$2-$\alpha$2 angle (right).

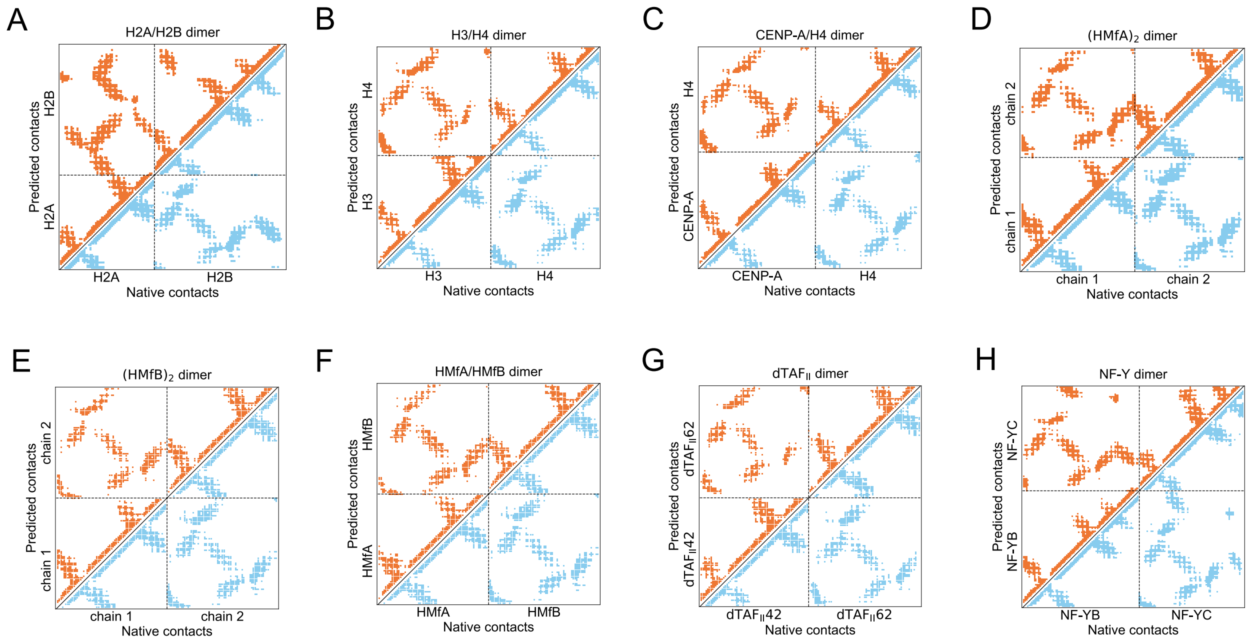


Figure S5: **Contact information of all the histone-like protein dimers in this study is accurately predicted by our simulations.** Contact maps are plotted for all the histone-like protein dimers in this study: (A) H2A/H2B; (B) H3/H4; (C) CENP-A/H4; (D) $HMfA_2$; (E) $HMfB_2$; (F) HMfA/HMfB; (G) $dTAF_{II}42/dTAF_{II}62$; (H) NF-YB/NF-YC dimer. Native and predicted contacts are represented in blue and orange, respectively. Predicted contacts for each dimer are computed using the structure of a simulation snapshot with the highest $Q$ value from ten annealing runs.

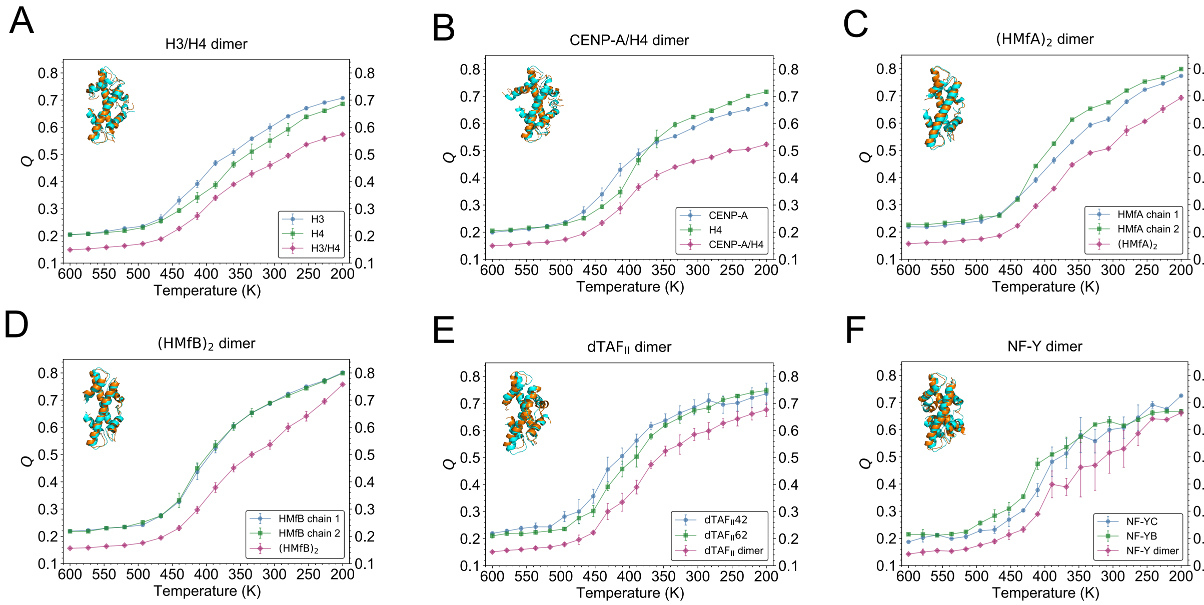


Figure S6: **Histone dimers and histone-like dimers fold upon binding.** $Q$ values for the folded monomer and dimer are shown as functions of the annealing temperature in AWSEM-MD simulations of H3/H4 (A), CENP-A/H4 (B), $HMfA_2$ (C), $HMfB_2$ (D), and histone-like dimers $dTAF_{II}42/dTAF_{II}62$ (E), NF-YB/NF-YC (F). Markers and error bars represent the mean values and standard deviations of $Q$ from ten independent simulation runs.

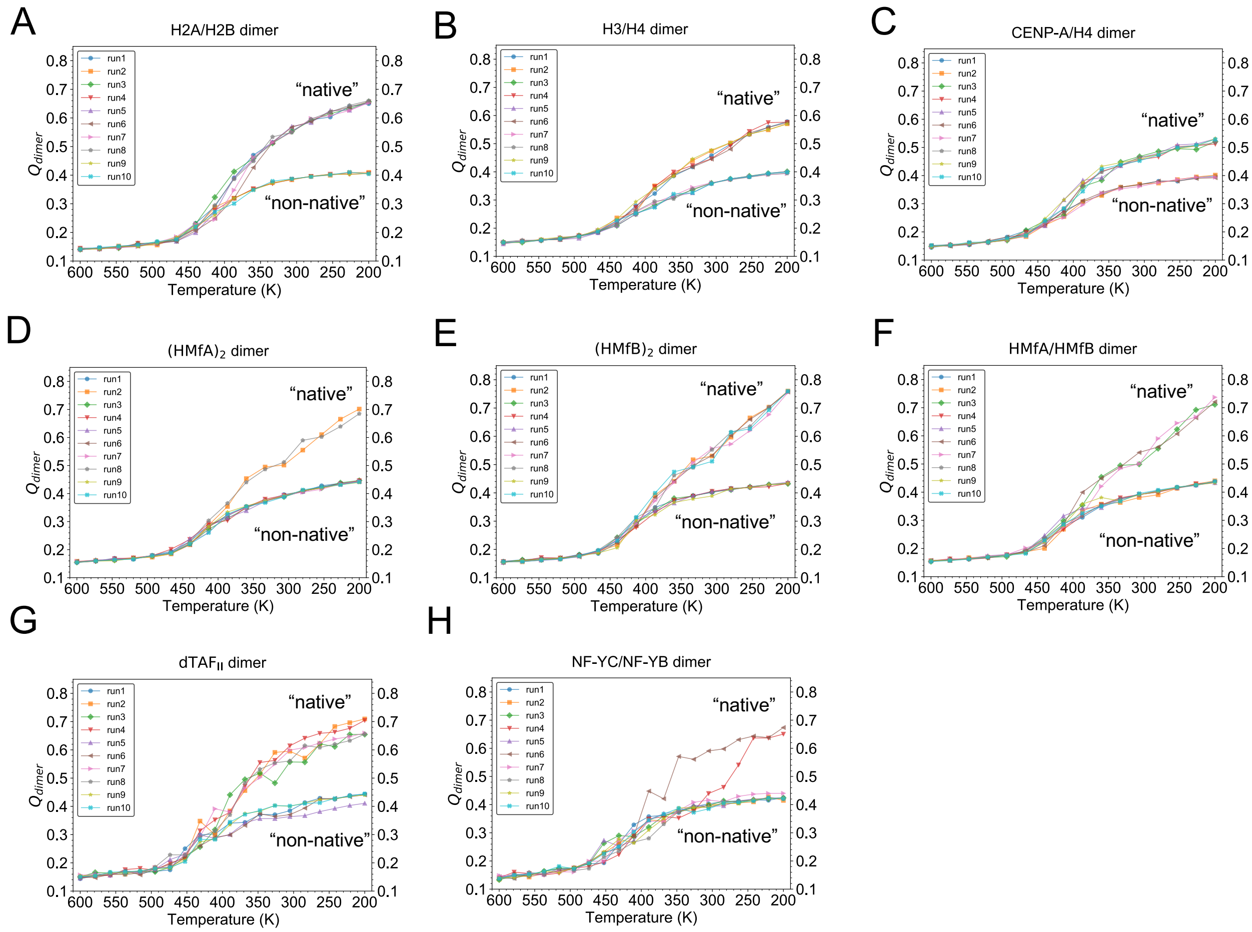


Figure S7: **Histone dimers and histone-like dimer proteins fold into native and non-native conformations.** $Q_{dimer}$ values are shown as functions of the annealing temperature for AWSEM-MD simulations of H2A/H2B (A), H3/H4 (B), CENP-A/H4 (C), $HMfA_2$ (D), $HMfB_2$ (E), HMfA/HMfB (F) and dimers $dTAF_{II}42/dTAF_{II}62$ (G), NF-YB/NF-YC (H). Ten individual simulation runs are represented in different colors and marker types. We categorize all the runs with a final $Q > 0.5$ as “native” and $Q < 0.5$ as “non-native”.

| Protein | H2A/H2B | | H3/H4 | |
|---|---|---|---|---|
| | Native | Non-native | Native | Non-native |
| PE_AWSEM | -568.25 ± 10.27 | -571.27 ± 7.74 | -546.48 ± 9.14 | -538.62 ± 15 |
| Protein | CenpA/H4 | | HmfA/HmfB | |
| | Native | Non-native | Native | Non-native |
| PE_AWSEM | -588.09 ± 12.41 | -587.2 ± 13.59 | -511.38 ± 12.66 | -504.82 ± 7.21 |
| Protein | (HmfA)_2 | | (HmfB)_2 | |
| | Native | Non-native | Native | Non-native |
| PE_AWSEM | -568.25 ± 10.27 | -571.27 ± 7.74 | -546.48 ± 9.14 | -538.62 ± 15 |
| Protein | dTAF_II42/dTAF_II62 | | NF-YB/NF-YC | |
| | Native | Non-native | Native | Non-native |
| PE_AWSEM | -588.09 ± 12.41 | -587.2 ± 13.59 | -511.38 ± 12.66 | -504.82 ± 7.21 |

Figure S8: **Potential energy from AWSEM-MD simulations shows both histone native and non-native conformations are energetically nearly degenerate.** The potential energy in AWSEM-MD includes the follow terms: $V_{backbone}$, $V_{contact}$, $V_{burial}$, $V_{HB}$, $V_{AM}$ and $V_{DSB}$ with details described in ref.[2]. Numbers in this table are in the unit of kcal/mol.

### S5 Supplemental NMR Results

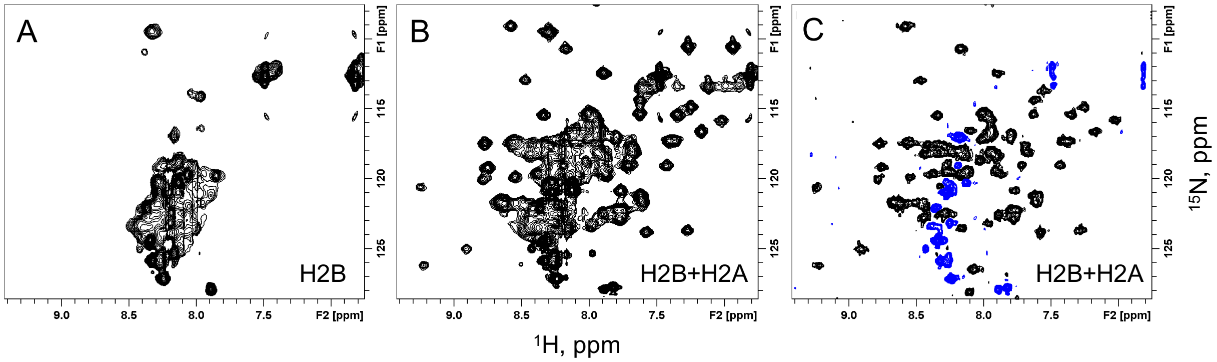


Figure S9: **NMR experiments of H2B upon complex formation** $^{1}$H-$^{15}$N SOFAST-HMQC spectra of $^{15}$N-labeled H2B alone (A) and in the presence of unlabeled H2A at an equimolar ratio (B). (C) Heteronuclear steady-state $^{15}$N-$^{1}$H NOE spectra recorded with amide proton presaturation for $^{15}$N-labeled H2B in the presence of unlabeled H2A at an equimolar ratio. In these spectra contours with positive intensities are colored black while negative intensities are blue.

### S6 Supplemental All-atom Simulation Results

Structures of predicted native, and non-native structure from AF2..

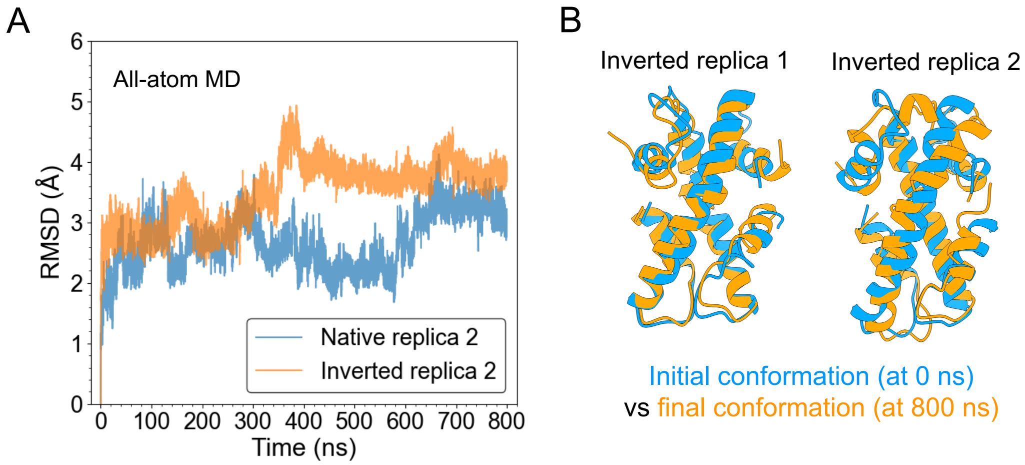


Figure S10: **The inverted H2A/H2B conformation is structurally stable in all-atom simulations** (A) Comparable stability of the inverted non-native conformation is consistently found to that of native structure in different simulation replicas. (B) The initial and final conformation (blue vs. orange) of the inverted structure display minor structural changes before and after 800-ns MD simulations.

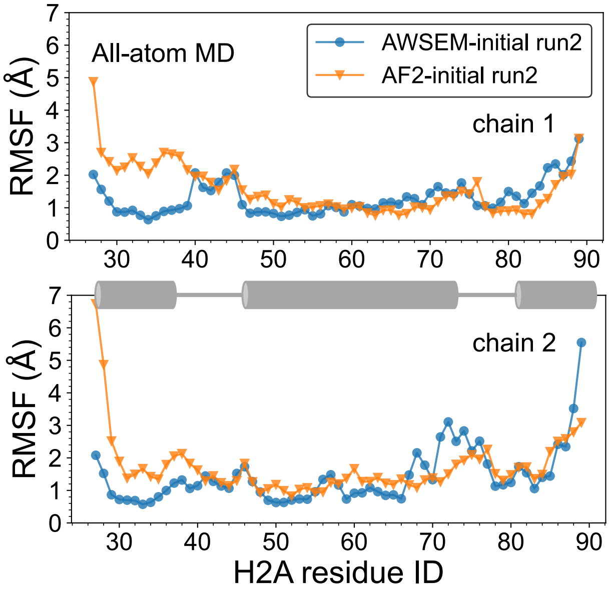


Figure S11: **The homo-complex structures of H2A/H2A predicted by AWSEM and AlphaFold2 have similar structural flexiblities.** The RMSF analysis of all-atom simulations exhibit similar atomic flexibilites of AWSEM- and AlphaFold2-predicted homo-complex structures in different simulation replicas. The RMSF of two chains are plotted separately where their helix regions are schemed by cartoon in the middle.

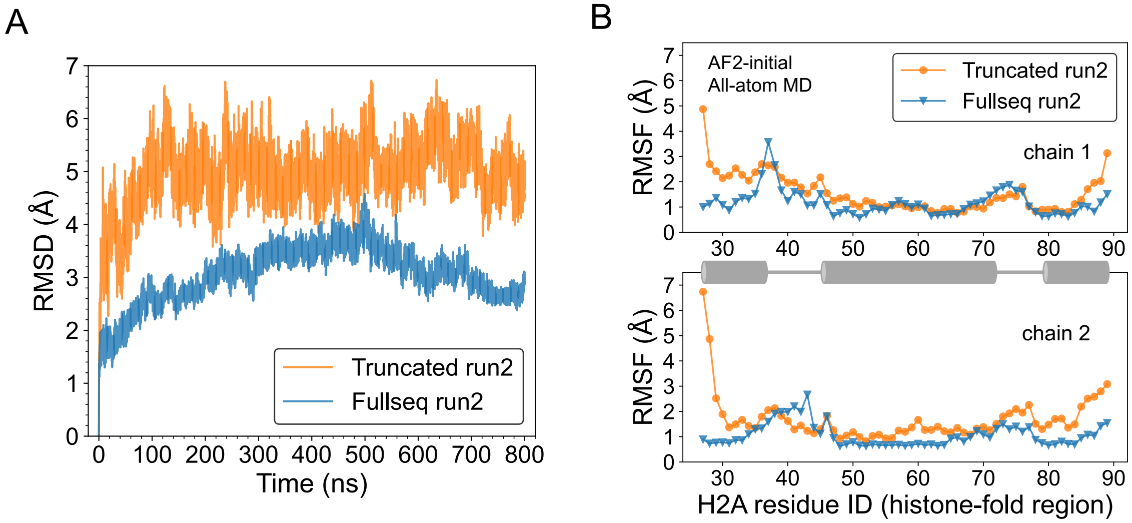


Figure S12: **Full-sequence histone homodimers show more stablity than that of only histone-fold region.** (A) The histone-fold region in full-sequence homodimers has less RMSD (blue) than that of histone-fold only structures (orange) in different simulation runs. (B) The RMSF analyses in different runs demonstrate that the two ending sessions are particulary flexible in trancated homodimers.

### S7 Discussion for Histone Folding/Binding Mechanism and Complex Structure Prediction

From the protein folding and binding theory perspective, a previous study by Levy *et al.* suggested that as in protein folding, the native topology of proteins is the major factor that determines their folding upon binding mechanism[17]. In the scenario of histones, the "hand-shake" geometry of histone-fold structure implies that histone may adopt an "induced-fit" mechanism model of protein-protein association[18]. Thus, it may not be surprising to realize structures like histone fold to have an induced-fit, or a coupled folding and binding mechanism. On the other hand, the unique sequence symmetry of histones with conserved hydrophobic residues lead the two participating monomers to fit with the induced formation but in two different ways, wherein the reserved hydrophobic residues play a dominant role. Interestingly, a previous experimental study found that H2A and H2B first rapidly recognize each other to form two intermediate bound states via weak hydrophobic interactions, and then rearrange to fold as the native structure[19]. This finding supports our hypothesis that the symmetrical hydrophobic interactions is essential for the induced-fit folding and binding process of histones. Another work by Zhou and coworkers showed that the foldability of a protein with native-reversed sequence depends on the protein size and location of its native hydrophobic core[20]. Here, in the histone binding example, we see that the conserved hydrophobic positions supports the N-C-terminal flipped binding conformations between two histones.

Protein structure prediction has been a long challenging problem since 1970s. Recently, breakthrough has been made through deep-learning based algorithms[21,22] and it is viewed as a stunning advance on solving the protein-folding problem. In this work, to predict homodimer structure of histones we applied the cutting-edge deep-learning based algorithm AlphaFold in ColabFold[23]. ColabFold is an online platform for protein folding and homo- and heteromer complex folding. Two models from AlphaFold2 were adpated: one is the

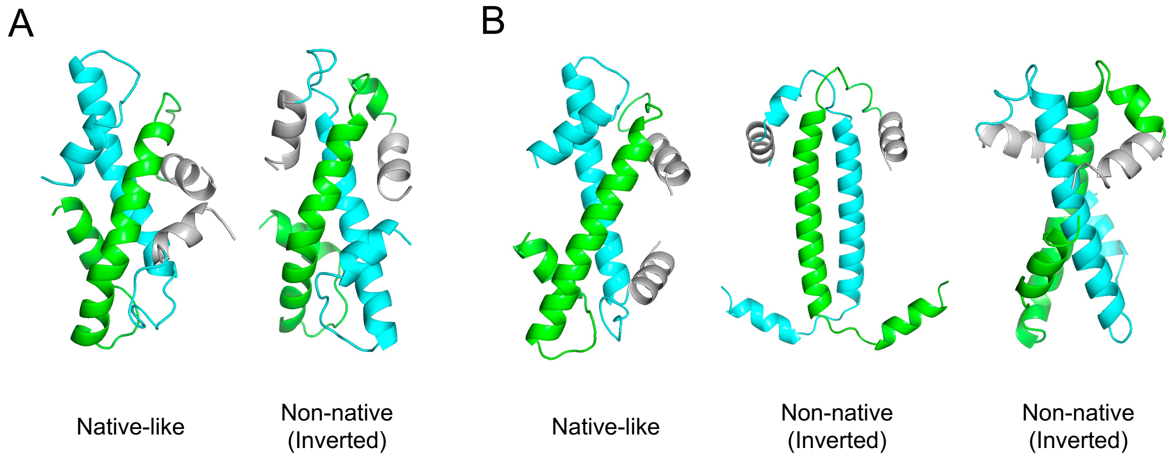


Figure S13: **Predicted structures of homo-complex of H2A/H2A by AWSEM and AlphaFold2.** Predicted strucrures of H2A/H2A by AWSEM (A) and AlphaFold2 (B) are shown, respectively. The two chains are colored in green and cyan while their $\alpha 1$ helixes in grey to help illustrate their native or non-native arrangements.

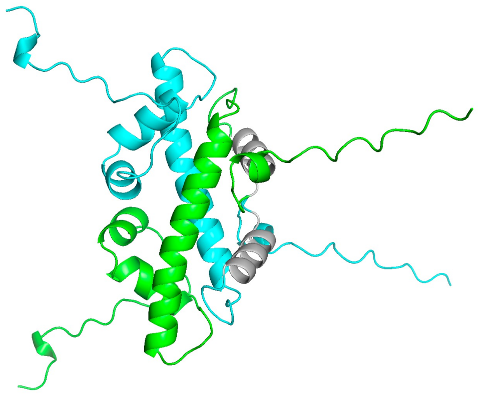

Figure S14: **AlphaFold2-predicted full-sequence homo-complex of H2A/H2A** The two chains are colored in green and cyan with their $\alpha 1$ helixes in grey. This prediction depicts a native-like arrangement of the histone fold structure.

original AlphaFold2 model[22] which was trained for monomeric protein folding; the other being the AlphaFold-Multimer[24] which is an AlphaFold2 model but trained specifically for multimeric inputs of known stoichiometry.

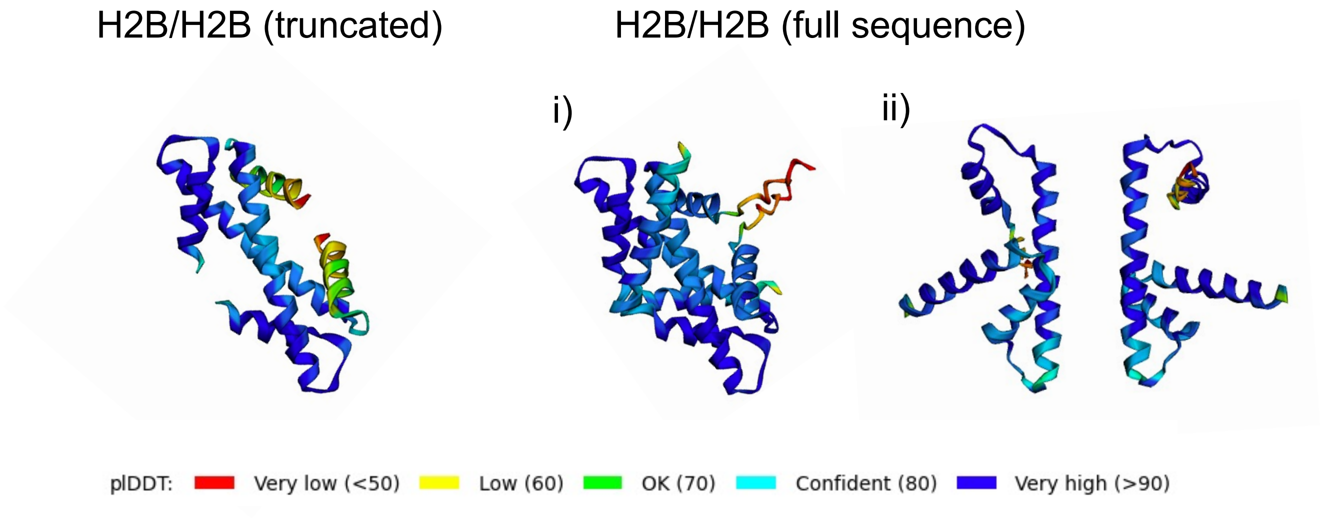


Figure S15: **AlphaFold2-predicted trancated and full-sequence homo-complex of H2B/H2B** The two chains are colored by the same scheme which is the plDDT confidence score provided by AF2. Both trancated and full-sequence H2B/H2B are predicted to have a native-like handshake structure while juxtaposing monomers are also found (full-sequence structure ii) which potentially indicates two non-interacting monomers.

Using ColabFold-adapted AF2 monomer model, we were able to predict complex for the H2A/H2A and got the native-like histone fold for H2A/H2A, while using AlphaFold-multimer, we obtained the inverted crossing way of histone-fold structure as what we found in AWSEM simulations (Figure S13). In total, the predictions of AF2 show outstanding consistency with that of AWSEM, which is mostly based on protein folding funnel energy landscape theory and coarse-grained MD simulations. The fact of different predictions out of two versions of AlphaFold approaches highlights the importance of training process in such methods. It is also possible that the AF-multimer is overtrained on available protein structures and have missed other possible states. One recent study indicates that modifying the multiple sequence alignment depth with stochastic subsampling may help generate alternative conformations[25], and another work points that optimising the multiple sequence alignment improves the precision of protein-protein interaction predictions[26].

As it is known, AlphaFold2 is an end-to-end protein structure prediction algorithm based on protein sequences and native protein structures, where the folding dynamics is completely absent. Together with our MD simulations results at both coarse-grained and all-atom resolutions, this work suggests that histone tails can increase the formation energy barrier of histone homo-dimerization likely by perturbing the structural fitting of histone-fold handshake core. Yet, once the homodimer has overcome the energy barrier, histone-fold structure may be formed and stay as stable.
